# A mechanistic, mortality-referenced biological age from routine blood tests: construction and temporal validation of IQN-BIOAGE-01

**DOI:** 10.64898/2026.09.29.755448

**Authors:** Igor Goryanin, Bob Damms, Irina Goryanin

## Abstract

**Background:** Blood-based biological-age clocks predict mortality well, but they are statistical composites. They do not say which physiological systems make a person biologically older, and they cannot say how an intervention would move the clock or whether moving it would change risk.

**Methods:** We built IQN-BIOAGE-01, a quantitative systems pharmacology (QSP) model of mortality-referenced biological age. A systemic senescent load follows saturating-removal stochastic dynamics and drives eight latent organ/hallmark axes, which are observed through 13 routine blood, blood-pressure and body-size measurements. Death is the first passage of the load over a threshold lowered by poor organ reserve, and biological age (BA_mech) is the age at which the reference population has the same 10-year risk. The model was calibrated on NHANES III (14,049 adults; 2,736 deaths in 20 years) and tested on a locked NHANES IV 1999–2010 hold-out (24,650 adults; 1,867 deaths). Two later versions were accepted or rejected by rules written before their validation results. We confronted the inflammation axis with three anti-inflammatory outcome trials, audited common implementations of the comparator clock, and locked a registration-ready external validation.

**Results:** On the hold-out, Harrell’s C was 0.842 for BA_mech, 0.842 for PhenoAge and 0.823 for chronological age (ΔC versus PhenoAge −0.0003, 95% CI −0.0026 to +0.0020). The hazard ratio per SD of the biological-age gap was 1.55 (1.48–1.62) for BA_mech and 1.45 (1.40–1.50) for PhenoAge. Risk ordering transferred across eras (calibration slope 1.01–1.03), but absolute 10-year risk was over-predicted (10.4% against 7.4% observed); life-table recalibration moved observed-to-expected from 0.72 to 0.81. The inflammation axis carried most of the gap. A joint re-estimate did not converge and a converged, biological-variation-anchored version narrowly failed its pre-written identifiability criterion, so v1.0 remains the locked primary. The outcome evidence available so far, chiefly top-line results from one anti-inflammatory trial, suggested that only a small fraction of a pharmacological CRP fall may carry the risk implied by the model’s cross-sectional structure (posterior median pass-through 0.22). Common software implementations of PhenoAge differed from the published formula by 0.7–2.0 years.

**Conclusions:** A mechanistic model can match a leading statistical blood clock in discrimination while adding an organ-level decomposition, explicit statements of what the data can and cannot identify, and a structure in which interventions can be simulated. Because inflammation dominates blood-based biological age while lowering CRP with drugs may carry little of the implied risk, a change in a blood clock should not be read as a change in mortality risk without outcome evidence. Validation is Tier 1 (internal and temporal, within one survey programme); the external test is locked and awaiting data access.

## Introduction

Biological-age clocks are now used as research endpoints and marketed to consumers, yet the best blood-based clocks remain regression composites. PhenoAge weights nine blood markers and chronological age by their association with mortality [1]. Such a score ranks risk well, but it cannot say which physiological system drives a high score, and it cannot simulate what an intervention would do.

Trials show why this matters. In CALERIE-2, two years of caloric restriction slowed blood-chemistry clocks [2, 3] and, modestly, the DNA-methylation pace measure DunedinPACE, but not DNA-methylation PhenoAge or GrimAge [4]. A clock built from the very markers an intervention changes will move whether or not risk has changed. Separating a real shift in risk from a change in the clock’s inputs requires a model of how the markers arise and how they relate to death.

Quantitative systems pharmacology (QSP) provides that structure. It represents physiology as state variables, links them to measurements through explicit observation equations, and links states to outcomes through a mechanism. For ageing, the saturating-removal model of senescent-cell turnover supplies such a mechanism: production rises with age, removal saturates, and first passage over a threshold reproduces the Gompertz law [5].

Here we describe IQN-BIOAGE-01, a QSP model of mortality-referenced biological age built on that core. We report its construction and calibration on individual-level data; its discrimination and calibration on a locked temporal hold-out against PhenoAge and chronological age; the decomposition of each person’s biological-age gap into organ/hallmark axes; the identifiability of its parameters and the pre-written rules that governed every model update; a test of whether its inflammation axis carries risk in the way anti-inflammatory outcome trials would require; an audit of the comparator clock’s implementations; and a locked, registration-ready external validation. Intervention and trial-design applications of the model are reported separately.

## Methods

### Data

Calibration used NHANES III (1988–94): 14,049 adults aged 20–84 years with all 13 markers, of whom 14,034 had mortality linkage and 2,736 died within 20 years. The temporal hold-out was NHANES IV 1999–2010: 24,650 adults, 1,867 deaths, median follow-up 9.4 years. Public-use files with linked mortality were obtained through the BioAge R package [2] and are not redistributed. The 13 markers are albumin, creatinine, glucose, C-reactive protein (CRP), lymphocyte percentage, mean corpuscular volume (MCV), red-cell distribution width (RDW), alkaline phosphatase (ALP), white-cell count (WBC), HbA1c, blood urea nitrogen (BUN), systolic blood pressure and body-mass index (BMI), together with age and sex. CRP values at the NHANES III assay floor (0.21 mg/dL) were treated as left-censored rather than imputed.

### Model structure

A single systemic senescent load X follows the saturating-removal stochastic model of Karin et al. [5], with t the age in years and time in days (Figure 1):

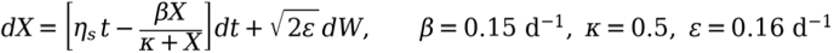

The removal constants β and κ come from the source. The sex-specific production slope η_s (men 0.001258, women 0.001176 per day per year) and the noise ε were refitted to NHANES III mortality after the source parameter set had been replayed against seven published anchors.

**Figure 1.**
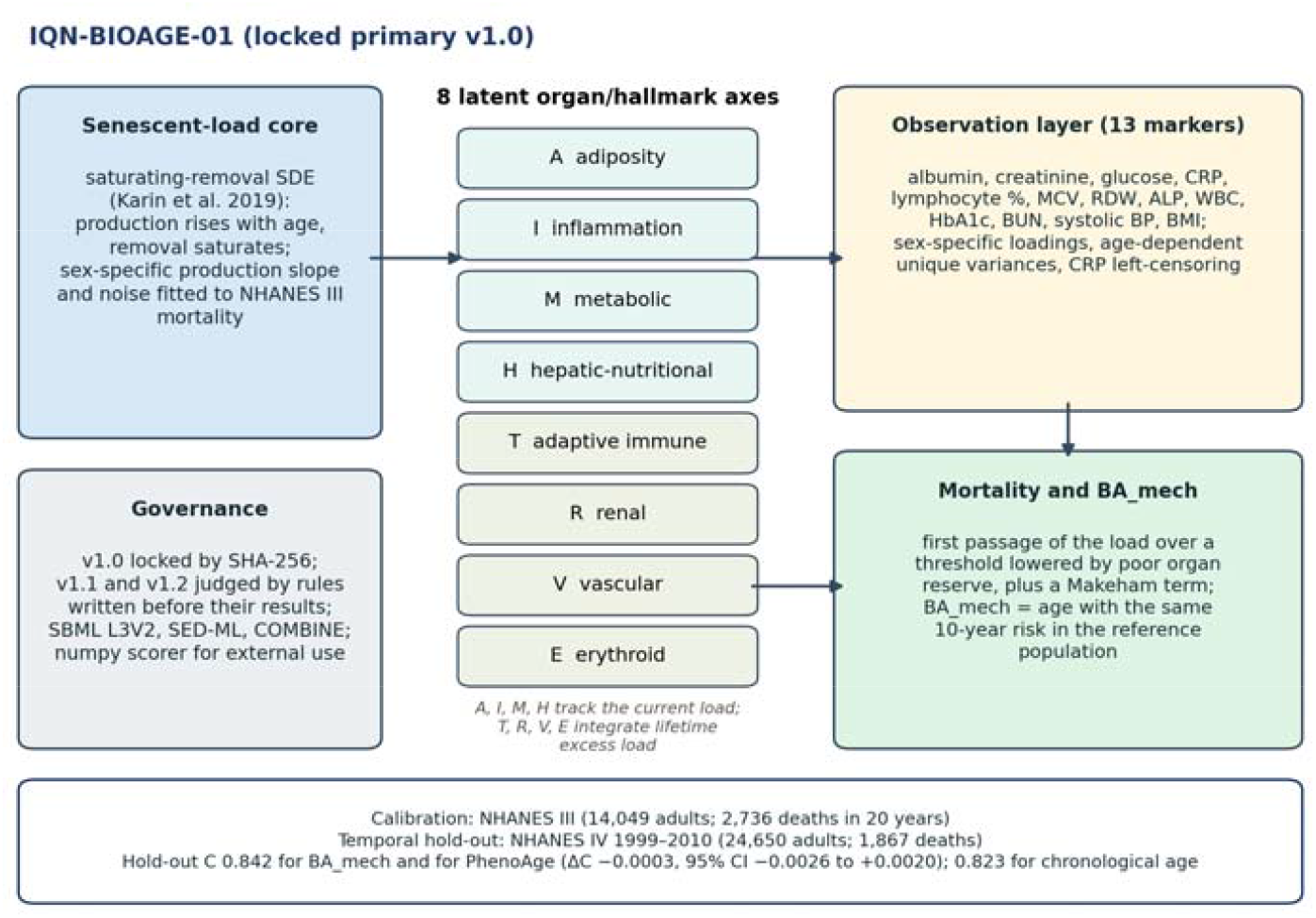
Structure of IQN-BIOAGE-01. A saturating-removal senescent-load core drives eight latent organ/hallmark axes; four reversible axes track the current load and four accumulating axes integrate lifetime excess load. The axes generate 13 routine markers through explicit observation equations and lower an individual first-passage mortality threshold. BA_mech is the age with the same 10-year risk in the reference population. The primary version (v1.0) is locked by SHA-256 hash, and later versions were judged by rules written before their validation results.

X drives eight latent axes: adiposity (A), inflammation (I), metabolic (M), hepatic-nutritional reserve (H), adaptive-immune decline (T), renal (R), vascular (V) and erythroid (E). The reversible axes A, I, M and H track the current load, whereas the accumulating axes T, R, V and E integrate lifetime excess load. Structural couplings carry effects between axes (adiposity to inflammation; inflammation and adiposity to metabolic; metabolic and adiposity to vascular; inflammation to erythroid), and person-level random intercepts and slopes give each axis its own level and pace. Each marker is an explicit function of the axes, with sex-specific intercepts and loadings and unique variances that grow with age as resilience is lost (critical age 130 years, within the 120–150-year range estimated by Pyrkov et al. [6]). Two residual correlations (MCV–RDW and lymphocyte–WBC) are included.

Death is the first passage of X over an individual threshold that falls as organ reserve worsens, plus a constant background (Makeham) hazard. First-passage survival is computed exactly on a one-day Markov-chain grid. BA_mech is the age at which the pooled reference population has the same 10-year risk as the person, found by inverting the reference risk curve on the complementary log-log scale:

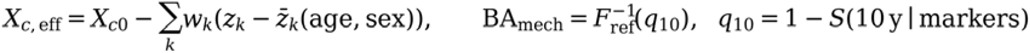

### Estimation: three model versions

Version 1.0 was estimated by staged maximum likelihood: 92 marker-model parameters, then 10 survival parameters, with unique variances free. Version 1.1 maximised the joint likelihood of markers and survival, with exact integration over censored CRP and Gauss–Hermite quadrature over axis uncertainty. Version 1.2 used the same joint likelihood fitted to full convergence (L-BFGS-B, then an active-set safeguarded Newton method, with Karush–Kuhn–Tucker conditions checked at bounds) and anchored the measurement model to published biological variation: each marker’s unique variance had to cover its published within-person variation plus desirable analytical imprecision (CVA = 0.5 × CVI) [7] and, for systolic blood pressure, a within-person SD of 8.2 mmHg [8]. This value was fixed for single-indicator axes (albumin, lymphocytes, blood pressure), which identifies them through reliability, and acted as a floor for the other markers except BMI. The two residual correlations were bounded to ±0.8, and two random slopes with no information were fixed.

### Validation metrics and recalibration

Discrimination was Harrell’s C for all-cause death, with paired percentile-bootstrap confidence intervals for differences (200 replicates in development; 1,000 are specified for the external test). Calibration was assessed as observed 10-year risk (1 − Kaplan–Meier) against predicted risk, the observed-to-expected ratio (O/E) and the Cox calibration slope of the log cumulative predicted hazard. Association was the hazard ratio per SD of the biological-age gap (BA_mech − age), adjusted for age and sex, with cause-specific Cox models for organ-matched deaths. For life-table recalibration, each person’s yearly hazard increments were scaled by the ratio of United Nations World Population Prospects 2022 single-year mortality (country, calendar year, age, sex) [9] to the NHANES III person-time reference; a secondary variant added participation multipliers for follow-up years 1–5 and 6–10 estimated in NHANES III. Neither used outcome data from the target cohort. PhenoAge was computed with the paper-faithful formula [1] throughout.

### Selection rules written before results

Every new version was judged by a rule written and time-stamped before its discrimination, calibration and identifiability results existed (v1.1 at 18:35 UTC and v1.2 at 20:51 UTC on 25 September 2026). Two process details are disclosed: the v1.1 rule was written while its fit was running, and the v1.2 identifiability criterion was coded after the fit log had shown which parameters sat at bounds but before the Hessian, identifiability and validation results existed. The v1.2 rule required all four of: convergence (Newton decrement below 1, positive-definite Hessian, KKT conditions at active bounds); a lower 95% bound of ΔC against v1.0 on NHANES IV above −0.002; an NHANES IV calibration slope within 0.8–1.2 after period recalibration; and no parameter non-identifiable in v1.2 that was identifiable in v1.0. Once inspected, NHANES IV was treated as a re-used hold-out, so later versions cannot claim an independent test on it.

### Identifiability and sensitivity

A parameter was classed as non-identifiable if it loaded above 0.3 on a near-null direction of the Hessian (eigenvalue below the maximum × 10^−9^); otherwise its relative standard error (RSE) classed it as well (below 20%), moderately (below 50%) or weakly identified. Survival parameters also had profile likelihoods, and a global sensitivity analysis (Latin hypercube, 3,000 samples per block, ±30%, Spearman correlations) ranked the drivers of each endpoint.

### Interventions as axis shifts, and a lifestyle replay

Interventions enter the model only as exogenous shifts of the axes or of the load dynamics, with first-order onset and offset; the calibrated backbone is not changed. As a face-validity check that uses no drug, caloric restriction was represented by shifts calibrated to CALERIE-2 weight (−10.4%) [10] and CRP (about −40%) [11] only, and its effect on blood-chemistry clocks was then predicted and compared with published results that were not used for calibration [2, 3].

### Does a fall in CRP carry risk? The inflammation pass-through

The backbone assumes that a fall in the inflammation axis lowers risk exactly as much as the cross-sectional structure implies. To test this, an exogenous shift on the inflammation axis, φ_I, was split into a risk-carrying part and a marker-only part by a pass-through fraction ψ (Equation 3); the backbone corresponds to ψ = 1. Because BA_mech is computed from the markers, its response to a CRP fall depends only weakly on ψ, whereas predicted risk depends on it strongly.

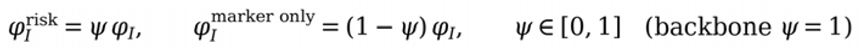

ψ was confronted with three outcome trials that report both a CRP change and a hard outcome. For each, the inflammation shift was fitted to the trial’s CRP change and the trial was replicated in a virtual cohort drawn from the calibrated model: CANTOS (canakinumab 150 mg; a 16,000-person virtual replica, mean age 61, hs-CRP at least 2 mg/L, 3.7 years) [12], colchicine after myocardial infarction [13], and ZEUS (ziltivekimab), for which only top-line results were available [14]. The CANTOS and ZEUS likelihoods were combined over a uniform prior on ψ. JUPITER was not used, because the model has no lipid axis and cannot represent a statin’s LDL-mediated benefit [15].

### MEASUREMENT properties for trials

To characterise BA_mech as a trial endpoint, we simulated parallel-arm trials of 24 months (12 months on treatment, then 12 months off) in 2,000 virtual participants per arm aged 45–70 years (BMI 27–35 kg/m^2^), comparing an untreated arm with an arm given the CALERIE-calibrated caloric-restriction shifts, with monthly blood panels. Each panel was the simulated latent state plus a persistent person-level component and transient noise drawn at every visit from published within-person biological variation with desirable analytical imprecision [7, 8] (1% day-to-day coefficient of variation for BMI), and every panel was scored with the locked scorer, as a real panel would be. The endpoint was the mean of the months 10–12 panels minus the mean of two baseline panels, compared by ANCOVA.

### Software, verification and the external-validation package

The reference implementation is in Python (JAX 0.4.35, NumPy, SciPy, statsmodels), with SBML Level 3 Version 2 models, SED-ML simulation descriptions and a COMBINE archive [16, 17]. The sex-specific SBML files replay the Python reference from age 20 to 85 within 0.018% (men) and 0.67% (women) of each marker’s age range, against a 1% tolerance, and libSBML reports no read or consistency errors. The locked external-validation package contains a stand-alone NumPy scorer, a statistical analysis plan (SAP 1.1), a UK Biobank field converter, life tables for 25 countries and automated tests. Every model and code file is hashed, and the runner re-verifies the hashes at start-up. Registration drafts follow the OSF Secondary Data Preregistration template [18].

The work was carried out with an AI-assisted modelling workflow that requires source replay before fitting, a register of claim warnings, explicit validation tiers and a closing model-status statement; independent verification agents that did not build the work audited each round (see Declarations).

## Results

### Discrimination: equal to PhenoAge, better than age

BA_mech discriminated all-cause mortality as well as PhenoAge and better than chronological age, both internally and on the locked hold-out (Table 1; Figure 2). On NHANES IV the difference from PhenoAge was −0.0003 (95% CI −0.0026 to +0.0020), within ±0.003 of zero, and the gain over chronological age was +0.0183 (+0.0155 to +0.0217).

**Table 1.** Discrimination and association with mortality (primary model v1.0).

| Quantity | NHANES III (calibration, 20-year follow-up) | NHANES IV 1999–2010 (locked hold-out) |
| --- | --- | --- |
| Adults / deaths | 14,034 / 2,736 | 24,650 / 1,867 |
| C, BA_mech | 0.860 | 0.842 |
| C, PhenoAge (paper-faithful, ln CRP) | 0.859 | 0.842 |
| C, chronological age | 0.846 | 0.823 |
| $\Delta C$, BA_mech – PhenoAge (95% CI) | $+0.0012$ ($-0.0002$ to $+0.0025$) | $-0.0003$ ($-0.0026$ to $+0.0020$) |
| $\Delta C$, BA_mech – age (95% CI) | $+0.0144$ ($+0.0124$ to $+0.0166$) | $+0.0183$ ($+0.0155$ to $+0.0217$) |
| HR per SD of gap, BA_mech (age/sex-adjusted) | 1.50 (1.45–1.56) | 1.55 (1.48–1.62) |
| HR per SD of gap, PhenoAge | 1.43 (1.39–1.48) | 1.45 (1.40–1.50) |
| Within-age-band C of the gap, BA_mech / PhenoAge | 0.635 / 0.632 | 0.636 / 0.635 |
| SD of the gap, BA_mech / PhenoAge (years) | 4.6 / 5.9 | 4.4 / 6.4 |

**Figure 2.**
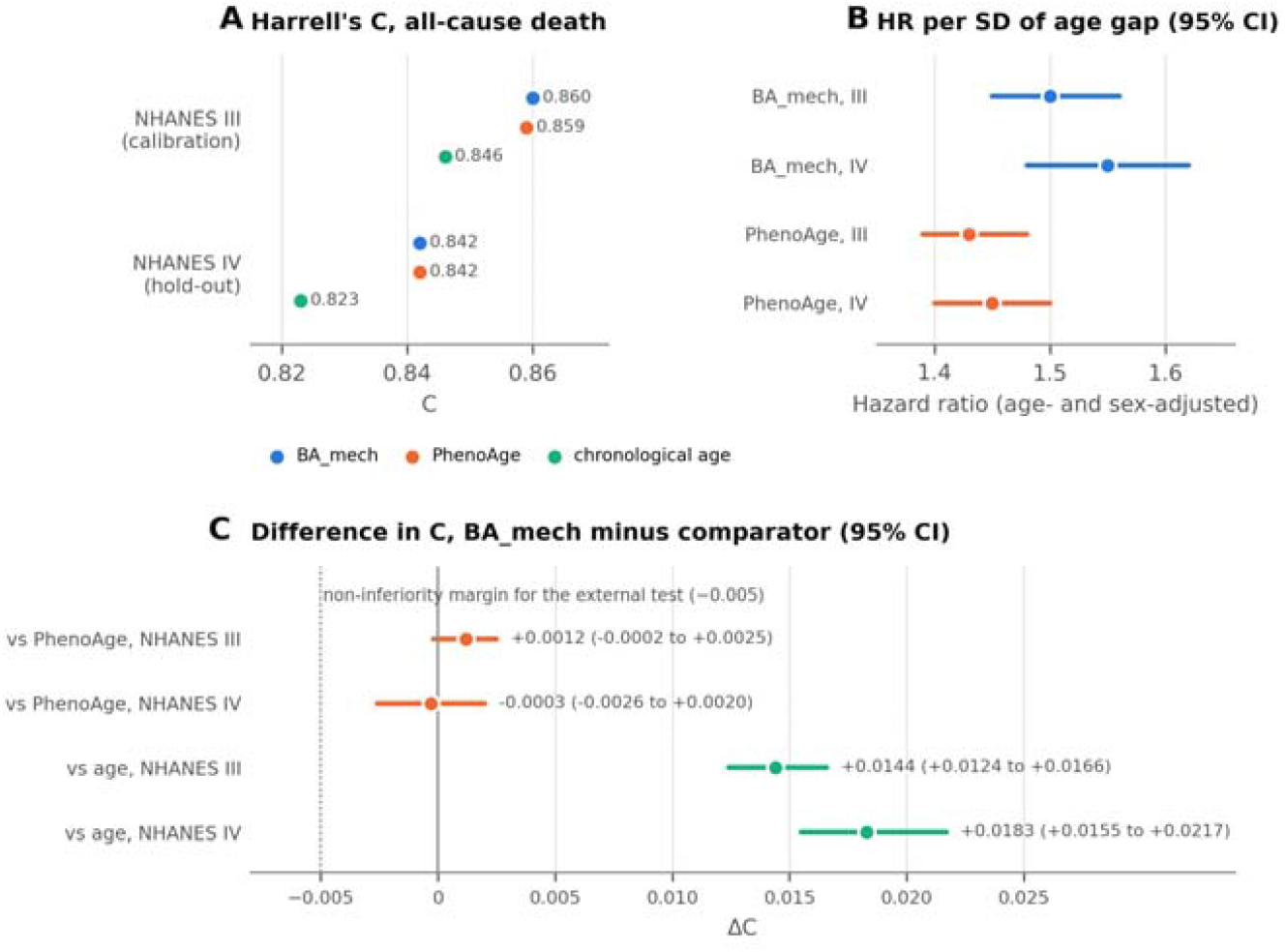
Discrimination and association with mortality. (A) Harrell’s C for all-cause death in the calibration cohort (NHANES III) and the locked temporal hold-out (NHANES IV 1999–2010) for BA_mech, PhenoAge (paper-faithful formula) and chronological age. (B) Hazard ratio per SD of the age- and sex-adjusted biological-age gap. (C) Differences in C with paired bootstrap 95% confidence intervals; the dotted line marks the non-inferiority margin pre-specified for the external test.

The two biological ages were highly correlated (r = 0.98), but their gaps from chronological age much less so (r = 0.74). BA_mech compresses the gap (SD 4.4 against 6.4 years) yet carries a stronger hazard ratio per SD, so each year of BA_mech gap carries more mortality risk than a year of PhenoAge gap.

### Absolute risk: right ordering, wrong level across eras

The model ranked risk correctly in both eras but over-predicted its level in the later one. Internally, mean predicted 10-year mortality was 10.3% against 9.9% observed. In NHANES IV it was 10.4% against 7.4% (Kaplan–Meier), yet the calibration slope stayed close to 1 (1.01, 95% CI 0.96–1.05; Figure 3A). Life tables closed about a third of the gap on the O/E scale (0.72 to 0.81), or 1.3 of the 3.0 percentage-point excess in mean predicted risk, the part due to falling population mortality between the eras: after period recalibration the mean prediction was 9.1% (O/E 0.81, slope 1.03, 0.98–1.07), and adding participation multipliers gave 8.8% (O/E 0.84). The remainder sat early in follow-up. After period recalibration, O/E was 0.70 in years 0–5 and 0.93 in years 5–10 in NHANES IV, and 0.76 then 1.10 in NHANES III (Figure 3B): people well enough to be examined in a survey die less often than the population over the next few years. Absolute risk should therefore be quoted only with its recalibration variant and its O/E; discrimination claims do not depend on it.

**Figure 3.**
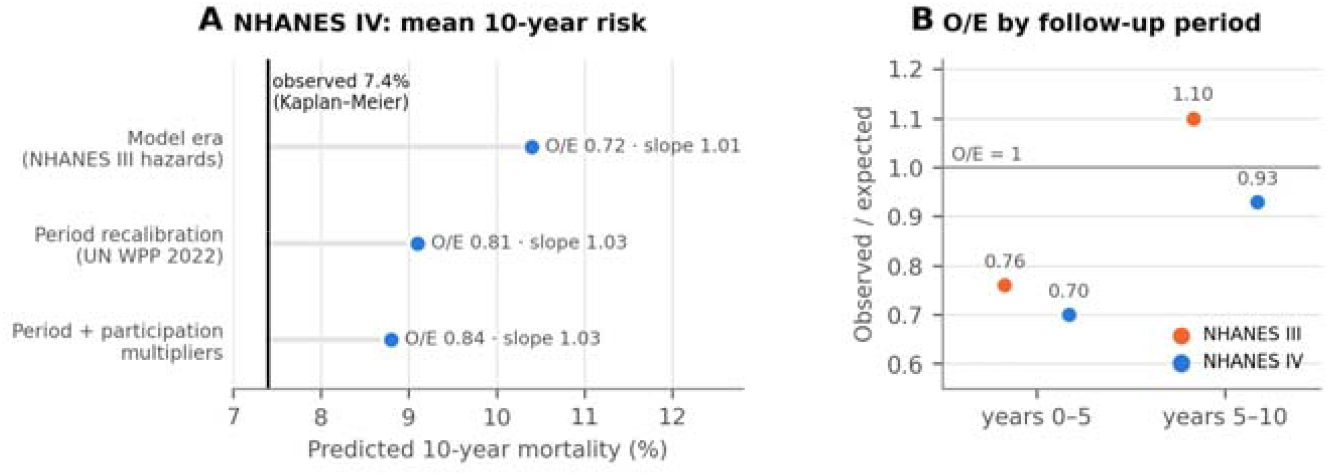
Absolute-risk calibration. **(A)** Mean predicted 10-year mortality in NHANES IV under the model-era hazards and two recalibration variants, with O/E and the Cox calibration slope; the vertical line marks observed mortality (Kaplan–Meier). (B) O/E by follow-up period after period recalibration in both cohorts; the early deficit of deaths is consistent with a healthy-participant effect.

### What drives the gap: inflammation first, organs only partly

Inflammation carried most of the biological-age gap, and the organ axes were organ-weighted rather than organ-specific. In NHANES IV the inflammation axis accounted for 108% of the gap variance, offset by the erythroid axis (−67%); the metabolic (17%) and vascular (13%) axes, the current senescent load (5.5%) and hepatic-nutritional reserve (5.2%) followed, and the adaptive-immune, renal and adiposity axes contributed less than 1% each (Figure 4A).

**Figure 4.**
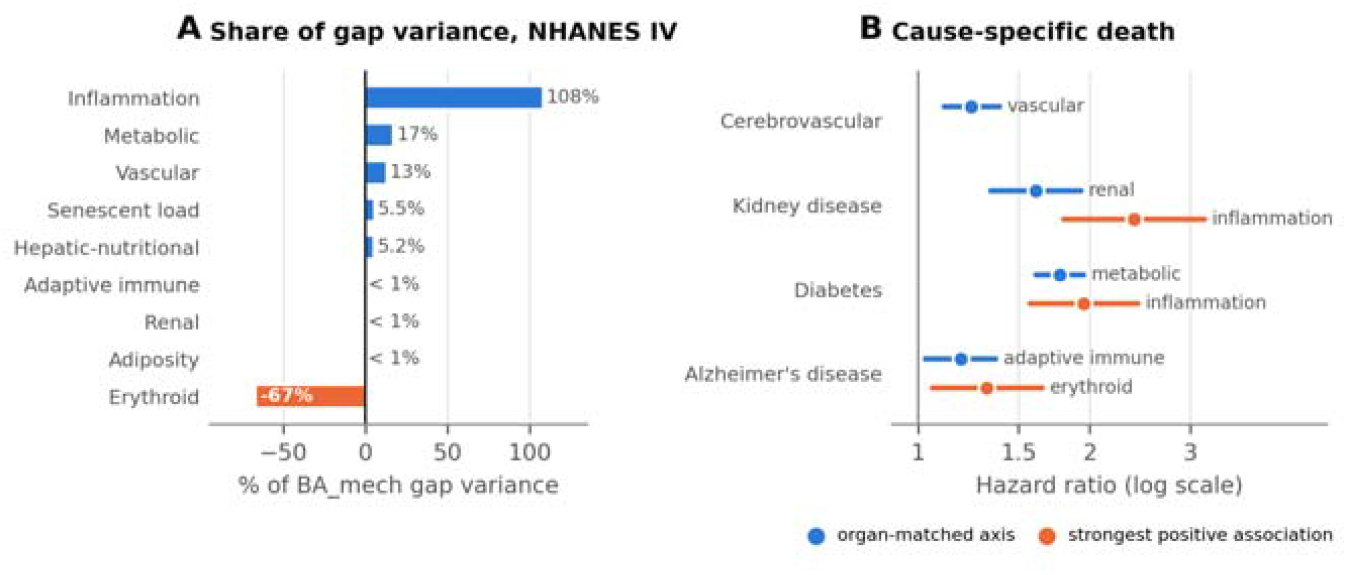
What drives the biological-age gap. (A) Share of the variance of the BA_mech gap attributable to each axis in NHANES IV; shares can exceed 100% because axes covary, and the erythroid axis offsets part of the inflammation signal. (B) Cause-specific mortality per SD of axis deviation (age- and sex-adjusted Cox models, NHANES IV): the organ-matched axis and, where different, the axis with the strongest positive association.

Organ-matched cause-specific associations were present but rarely the strongest (Figure 4B). Only for cerebrovascular death was the matched (vascular) axis the top predictor (HR per SD 1.24, 1.11– 1.39). For kidney-disease death the renal axis gave 1.61 (1.34–1.93) but inflammation 2.39 (1.80– 3.18); for diabetes death the metabolic axis gave 1.77 (1.61–1.95) and inflammation 1.95 (1.57– 2.43). Adiposity was inversely related to Alzheimer’s disease death (HR 0.60, 0.45–0.79), consistent with weight loss before diagnosis.

Extreme deviations tracked risk, but mostly through the total gap. Counting seven axes (adiposity excluded) with a one-sided z above 1.5 against the model’s age- and sex-specific distribution, capped at 3, 14.5% of adults had at least one extreme axis, and mortality rose with the count after adjustment for age and sex (HR per axis 1.70, 1.57–1.85). Adjusted also for the biological-age gap, the hazard ratio fell to 1.09 (0.98–1.20), and crude death rates were not monotone (7.0, 12.2, 13.7 and 8.7 per 1,000 person-years for 0, 1, 2 and 3 or more axes). Under the definition pre-specified for the external test (all eight axes, |z| above 1.5, cohort-standardised) the age- and sex-adjusted hazard ratio was 1.24 (1.20–1.28). Cross-axis correlations did not rise with age, so a single shared driver of all axes is not supported by these cross-sectional data. Organ “ages” in years were attempted but are not reported, because only four of the eight axes change monotonically enough with age to be mapped to years.

### Identifiability, and three versions judged by rules

The primary model is well identified where it matters for BA_mech. In v1.0, 79 of 92 marker-model parameters were well identified, 5 moderately and 8 not at all; the 8 were mostly variance splits of axes measured by a single marker (albumin, lymphocytes, blood pressure), plus the MCV loading on the erythroid axis and the BMI unique variance. In the survival block, profile-likelihood 95% intervals were 1.47–2.01 for the inflammation weight, 1.61–2.93 for metabolic, 0.18–0.35 for vascular and 55.3–80.5 for erythroid; the adiposity, adaptive-immune and renal weights were compatible with zero. Global sensitivity analysis showed that the model’s predicted response of BA_mech to weight-loss interventions such as caloric restriction depends mainly on the inflammation and erythroid weights and on one confounded coupling pair (inflammation-to-erythroid coupling against the RDW loading on inflammation, r = −0.87).

Two attempts to improve the model were rejected by the rules written before their results (Table 2). Version 1.1 improved the log-likelihood but did not converge. Version 1.2 met the convergence, discrimination and calibration criteria (NHANES IV slope 1.003) but failed the identifiability criterion: the Makeham background hazard, fitted at 2.7 × 10^−5^ per year, lay on a near-null Hessian direction with eigenvalue 0.138, just below the tolerance of 0.153. The failure is marginal, and an independent verification showed that other reasonable readings, such as a relative-SE judgement for this parameter as used in v1.0, a scale-invariant Hessian or a slightly smaller tolerance, would have passed v1.2. The rule was written first, so it was applied as written: v1.0 stays primary, v1.1 secondary, and v1.2 enters the external plan only as an exploratory comparator. Anchoring to biological variation also changed the survival weights materially (in v1.2 the inflammation weight was 0.81, Wald 95% CI 0.67–0.98, and the erythroid weight 24.2, 20.8–28.2) while leaving discrimination almost unchanged (BA_mech correlation between versions 0.99), so these weights should be read as scale-dependent rather than as physiological constants.

**Table 2.** Model versions and their pre-written selection rules.

| Version | Change in NHANES III log-likelihood | Convergence | NHANES IV C ($\Delta C$ vs v1.0, 95% CI) | Identifiability | Decision |
| --- | --- | --- | --- | --- | --- |
| v1.0, staged | Reference | Stage-wise | 0.8416 | 8 of 92 non-identifiable | Primary, locked |
| v1.1, joint | +52.4 (markers 7.6, survival 44.8) | Not converged: Newton decrement 1.73; variances drifting to bounds | 0.8442 (+0.0026, +0.0015 to +0.0037) | Weakly identified splits drift | Secondary, locked |
| v1.2, joint, anchored to biological variation | +26.9 (markers –18.1, survival +45.0) | Converged: decrement 0.007; positive-definite Hessian; KKT holds | 0.8445 (+0.0029, +0.0019 to +0.0040) | 6 of the 8 resolved; Makeham term newly on a near-null direction | Not adopted; exploratory comparator |

NHANES IV has now informed three selection decisions and no longer counts as an independent test, which is the main reason the external validation below matters.

### A lifestyle replay

Calibrated only on CALERIE-2 weight and CRP, the model predicted that caloric restriction slows blood-chemistry PhenoAge by 0.81 years per year relative to ad libitum intake. The observed values, which were not used for calibration, were 0.66 years per year for blood PhenoAge [2] and 0.60 (0.21–0.99) years per year for Klemera–Doubal biological age [3, 19]. The model thus reproduced the direction and approximate size of a published clock response from marker-level inputs alone.

### A fall in CRP may carry little risk

With full pass-through (ψ = 1), the model’s CANTOS replica gave an all-cause mortality hazard ratio of 0.92, inside the observed 0.94 (0.83–1.06) [12]; this trial is consistent with any ψ from 0 to 1 and matched its point estimate at ψ of about 0.77. For colchicine after myocardial infarction, the model gave 0.94 against an observed relative risk of 0.97 (0.85–1.11) [13]. In ZEUS, in which ziltivekimab lowered hs-CRP but did not reduce major adverse cardiovascular events (hazard ratio 0.99, 0.88– 1.11), the model at ψ = 1 predicted 0.80 (Figure 5B) [14]. Combining the CANTOS and ZEUS likelihoods over a uniform prior gave a posterior median ψ of about 0.22, with about 56% of the mass below 0.25. The estimate is fragile: the reweighted posterior rests on an effective sample of about eight draws, it uses a cardiovascular rather than an all-cause endpoint for ZEUS, and the CRP reduction assumed for ZEUS must be checked against the full report.

**Figure 5.**
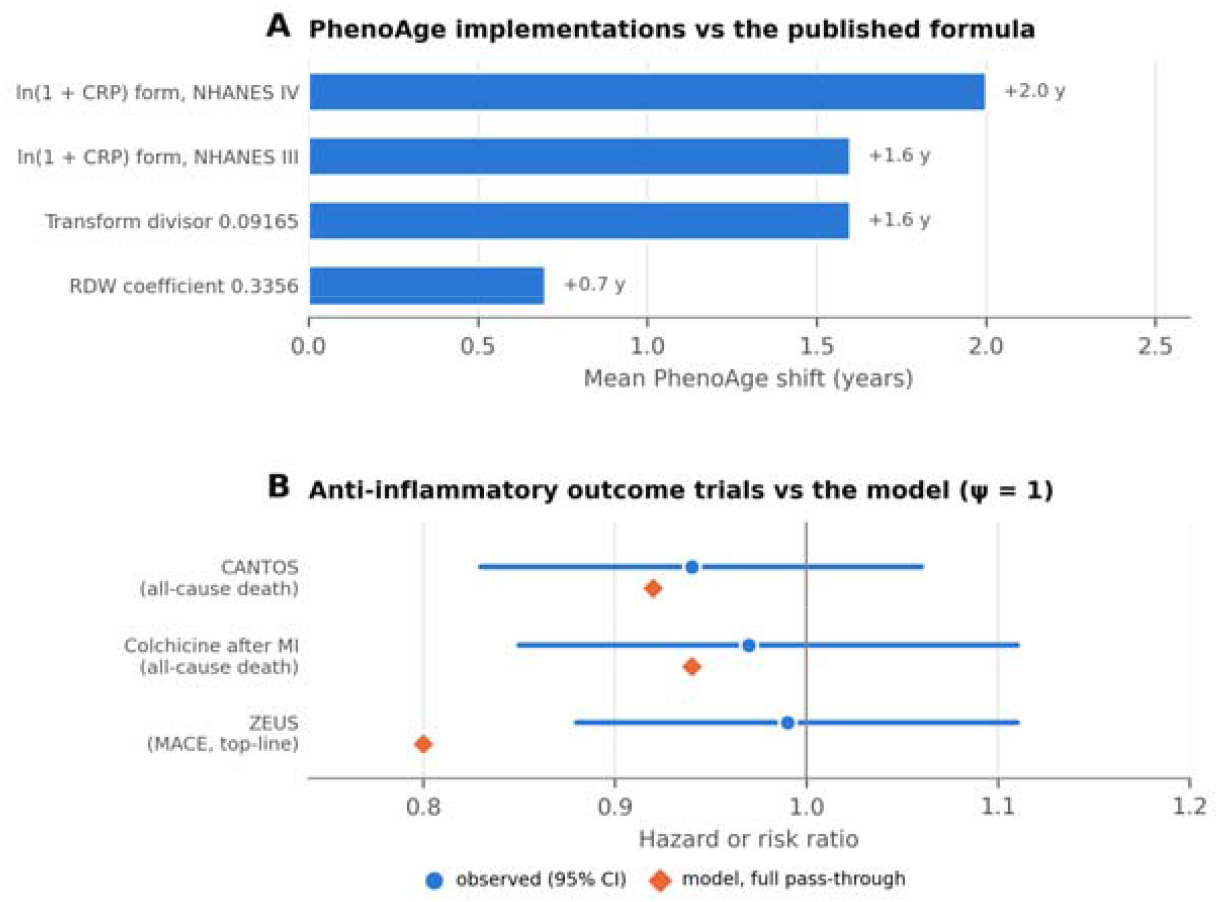
Measurement choices and the inflammation pass-through. (A) Mean shift of common PhenoAge implementations relative to the published formula: the ln(1 + CRP) form used by one widely used package, a transform divisor quoted in secondary sources (0.09165 instead of 0.090165) and an RDW coefficient in one open library (0.3356 instead of 0.3306). (B) Three anti-inflammatory outcome trials: observed hazard or risk ratio with 95% CI (circles) and the model’s prediction with full pass-through of the CRP fall into risk (ψ = 1; diamonds). ZEUS reports major adverse cardiovascular events from top-line results. The model at ψ = 1 over-predicts the benefit where CRP falls most.

The consequence for the model is specific. Inflammation dominates the biological-age gap (Figure 4A), and the backbone treats a fall in the inflammation axis as a proportional fall in risk. If ψ is low, an intervention that lowers CRP will lower BA_mech, and PhenoAge, by almost the same amount whatever ψ is, while lowering risk much less. A blood-clock change produced mainly through CRP should therefore not be interpreted as a change in mortality risk without outcome evidence.

### Measurement choices can be as large as treatment effects

PhenoAge is not one formula. Building ln(1 + CRP) instead of ln CRP, as one widely used package does, shifted PhenoAge by +1.6 years on average in NHANES III and +2.0 years in NHANES IV; an RDW coefficient of 0.3356 instead of the published 0.3306 in one open library added +0.7 years [20]; and a transform divisor of 0.09165 quoted in secondary sources, instead of 0.090165, added +1.6 years (Figure 5A). These variants barely changed ranking (the paper-faithful and ln(1 + CRP) versions gave the same C of 0.842 in NHANES IV), so discrimination studies are unaffected, but levels and changes are. The ln(1 + CRP) form also blunts the response to CRP: at the median CRP (0.2 mg/dL), a one-third fall lowers paper-faithful PhenoAge by about 0.43 years but the ln(1 + CRP) version by about 0.06 years, roughly seven-fold less (about two-fold less at 1 mg/dL). A trial that lowers CRP would report a smaller benefit with that implementation. Every PhenoAge result in years should name its implementation.

### BA_mech as a trial endpoint

In the simulated trials, BA_mech in the untreated arm rose by 0.97 years over 12 months and by 2.03 years over 24 months, close to one year per calendar year, as a mortality-referenced age should. The residual SD of a baseline-adjusted 12-month between-arm contrast was 2.26 years. The 95% minimal detectable change for an individual, between a two-panel baseline and a three-panel follow-up, was 4.4 years, larger than any treatment effect predicted in these simulations, so BA_mech is a group-level endpoint rather than a personal monitoring tool. Sampling density substituted for participants: for an effect of the size the model predicts for caloric restriction (1.32 years at 12 months), power at 50 evaluable participants per arm (two-sided 5%) was 0.83 with three-panel means from monthly sampling, 0.71 with quarterly panels and 0.61 with single panels at months 0 and 12.

### A locked, registration-ready external test

No external validation has been run yet; the test is prepared so that every analytic choice is fixed before external data are seen. The model files, scorer and analysis runner were locked by SHA-256 hash on 25 September 2026 (20:36:52 UTC) and re-verified after the v1.2 decision (21:42:58 UTC) with no mismatches. The stand-alone scorer reproduced the development pipeline to a maximum 10-year-risk difference of 6.9 × 10^−15^ in NHANES IV and a maximum BA_mech difference of 2.3 × 10?8 years. On 59,980 people simulated from the model itself (5,645 deaths), predicted risk was calibrated (O/E 0.994; slope 0.985, 0.953–1.017). Six automated tests pass, including the full runner on a synthetic cohort with missing markers. The pre-specified hypotheses and the wording allowed for each outcome are in Table 3.

**Table 3.** Pre-specified hypotheses for the external test (SAP 1.1) and the NHANES IV software run. NHANES IV was used during development, so this run tests the software and decision logic, not the model.

| Hypothesis | Decision rule | NHANES IV software run | Wording if met |
| --- | --- | --- | --- |
| H1: BA_mech discriminates death better than age | Lower 95% bootstrap bound of $\Delta C$ above 0 (one-sided $\alpha$ 0.025) | $\Delta C$ +0.018 (0.015 to 0.022) | Discriminates beyond age |
| H2: non-inferior to PhenoAge (tested only if H1 holds) | Lower bound of $\Delta C$ above -0.005 | $\Delta C$ -0.0003 (-0.0022 to 0.0019) | With H1 and H3: "externally validated for discrimination and risk ordering (Tier 2)" |
| H2b: superior to PhenoAge (tested only if H2 holds) | Lower bound of $\Delta C$ above 0 | Not met | Superiority claim allowed only if met |
| H3: risk ordering calibrated after period recalibration | Cox slope CI includes 1, or estimate within 0.8–1.2 | Slope 1.03 (0.98 to 1.07); O/E 0.81 | If not met, discrimination may stand but absolute risk is not transportable |

The software run used the locked runner on 24,646 adults (1,867 deaths) with its own bootstrap, so its intervals differ slightly from Table 1. The non-inferiority margin (0.005) is about 2.5 times the ΔC confidence half-width seen in NHANES IV, and a UK Biobank analysis would have roughly 20 times as many deaths. If H1 is not met, the pre-agreed wording is “not validated in the cohort”, with no refitting on the validation cohort. UK Biobank has paused new applications and intends to reopen them in late 2026 [21]; an application draft and an OSF pre-registration draft are ready, and any other cohort with all 13 markers and mortality follow-up can be scored with the same runner. The model is not validated with missing markers.

## Discussion

A mechanistic model of biological age ranked mortality risk as well as PhenoAge on a temporal hold-out and better than chronological age. Equal discrimination was the realistic ceiling: both scores read an almost identical blood panel, both were trained on NHANES III mortality and both were tested on the same NHANES IV adults. The mechanistic structure did not cost accuracy, and what it adds is different in kind: a decomposition of each person’s gap into axes, a count of extreme axes that flags multi-system deviation, explicit statements of which parameters the data can and cannot identify, and a structure in which interventions enter as explicit axis shifts.

The decomposition is the model’s most useful and most sobering output. Inflammation carries most of the biological-age gap, and the organ axes are organ-weighted rather than organ-specific. At the same time, the outcome-trial evidence available so far, chiefly the top-line ZEUS result, with colchicine pointing the same way and CANTOS compatible with any ψ, suggests that lowering CRP pharmacologically may carry only a small part of the risk that the cross-sectional association between CRP and death implies. Both statements may be true at once: inflammation can mark risk without being the lever that removes it. For blood-based clocks generally, this means that an intervention can move the clock largely through CRP without changing mortality to the extent the clock implies. Because BA_mech’s response to a CRP fall barely depends on ψ, no biomarker trial of this kind can settle ψ; outcome-anchored or genetic designs are needed.

In the model, blood-chemistry biological age behaves as a state as well as a trajectory. The reversible axes track the current load, which is consistent with the CALERIE pattern, in which caloric restriction moved blood-chemistry clocks [2, 3] but not DNA-methylation PhenoAge or GrimAge [4], and with the fast, recovering fluctuations of blood-marker state described in longitudinal data [6]. Trials with this endpoint should therefore make the between-arm contrast the primary estimand, sample densely, and include an off-treatment period if persistence matters; individual-level claims are not supported by a minimal detectable change of 4.4 years.

Absolute risk is era-specific. About a third of the over-prediction in NHANES IV (on the O/E scale) is the secular fall in mortality, which life tables correct; most of the rest is an early deficit of deaths among people well enough to be examined. Relative ranking transferred well. Absolute risk and BA_mech levels should be recalibrated in each new population and reported with their recalibration variant and O/E.

Rules written before results cost something and bought something. They rejected a non-converged joint fit and a converged version whose Makeham term lay on a near-null direction by a narrow margin, and they stopped us from adopting whichever version looked best on NHANES IV after seeing it. The price is that NHANES IV has now informed three decisions. Measurement choices matter as much: PhenoAge implementations differ by 0.7–2.0 years, comparable to many reported intervention effects, and weight loss can lower creatinine through lean-mass loss without any change in kidney function. Clock implementations, and a creatinine-independent renal marker such as cystatin C, should be specified before any trial reports a change in biological age.

The structure also limits some predictions. The model under-predicts the fall in blood pressure with weight loss (about −1.8 against about −9 mmHg per 10% weight loss in trials [22]), because the adiposity-to-vascular coupling was learned from variation between people and understates the within-person causal effect; a within-person pathway calibrated on weight-loss trials is planned as a new version judged by a pre-written rule.

## Limitations

Validation is Tier 1: all evidence comes from one survey programme, NHANES IV has informed three decisions, and no external cohort has been scored. Treating the axis-to-threshold link as causal is what allows interventions to be represented; it rests on cross-sectional associations, and the blood-pressure misfit and the low pass-through estimate show how within-person and pharmacological effects can differ. Several parameters are non-identifiable or weakly identified, and the v1.2 decision turned on an eigenvalue close to its tolerance. ψ is not identified by the available trials and rests on top-line results for one of them. CRP and albumin assays changed between NHANES III and IV, and marker harmonisation remains a source of error in any new cohort. The model uses 13 routine measurements at ages 20–84 years, includes no DNA methylation, functional measures or imaging, and is not validated with missing markers. The model, code and analyses were built with AI assistance and audited by independent AI verification agents; expert human review is required before any use beyond research.

## Conclusion

A mechanistic, mortality-referenced model of biological age built from 13 routine measurements matches PhenoAge in discrimination while decomposing each person’s gap into organ/hallmark axes and stating which of its parameters the data can identify. Inflammation dominates that gap, but the outcome evidence available so far, chiefly top-line ZEUS results, suggests that lowering CRP with drugs may carry little of the implied risk, so blood-clock changes need outcome evidence before they are read as changes in risk. Until an external cohort has been scored with the locked package, BA_mech is a Tier 1 research tool, and its absolute risks need local recalibration.

## Declarations

### Ethics

This is a secondary analysis of de-identified public-use NHANES files and simulated data. No participants were recruited. The UK Biobank analysis requires an approved application.

### Use of generative AI

Large language model agents executed parts of the modelling workflow, wrote and ran code, produced analyses and figures, and assisted in drafting this manuscript, under the direction of the author(s) IQANOVA Council (www.iqanova.org/aiqsp). Each round was audited by independent verification agents that had not produced the work. The author(s) checked the reported numbers against the source files and take full responsibility for the content. No AI system is an author.

### Author contributions

IG run the pipeline, BD edited the manuscript IVG provide admin support and funding. AI assistance is described above and is not an author contribution.

### Data and code availability

NHANES III and IV public-use files with linked mortality were obtained through the BioAge R package [2] and are not redistributed. United Nations World Population Prospects life tables are public [9]. The locked external-validation package (scorer, SAP 1.1, converters, life tables and tests) is identified by SHA-256 hashes. The SAP will be registered on OSF before any external data are accessed.

### Reporting

The prediction-model elements are to be reported against TRIPOD+AI [23]

